# Rat orbitofrontal ensemble activity contains a multiplexed but value-invariant representation of task structure in an odor sequence task

**DOI:** 10.1101/507376

**Authors:** Jingfeng Zhou, Matthew P.H. Gardner, Thomas A. Stalnaker, Seth J. Ramus, Andrew M. Wikenheiser, Yael Niv, Geoffrey Schoenbaum

**Affiliations:** Intramural Research Program of the National Institute on Drug Abuse, Baltimore MD; Bowdoin College, Bowdoin ME; Princeton Neuroscience Institute and Department of Psychology, Princeton University, Princeton NJ; Department of Anatomy and Neurobiology, Maryland School of Medicine, Baltimore MD; Department of Neuroscience, Johns Hopkins School of Medicine, Baltimore MD

**Keywords:** orbitofrontal, cognitive map, single unit, rat, expected value

## Abstract

The orbitofrontal cortex (OFC) has long been implicated in signaling information about expected outcomes to facilitate adaptive or flexible behavior. Current proposals focus on signaling of expected reward values versus the representation of a value-agnostic cognitive map of the task. While often suggested as mutually exclusive, these alternatives may represent two extreme ends of a continuum determined by the complexity of the environment and the subjects’ experience in it. As learning proceeds, an initial, detailed cognitive map might be acquired, based largely on external information. With more experience, this hypothesized map can then be tailored to include relevant abstract hidden cognitive constructs. This might default to expected values in situations where other attributes are minimized or largely irrelevant, whereas in richer tasks, a more detailed structure might continue to be represented, at least where relevant to behavior, and possibly alongside value. Here we sought to arbitrate between these options by recording single unit activity from the OFC in rats navigating an odor sequence task analogous to a spatial maze. The odor sequences provided a clearly mappable state space, with 24 unique “positions” defined by sensory information, likelihood of reward, or both. Consistent with the hypothesis that the OFC represents a cognitive map tailored to the subjects’ intentions or plans, we found a close correspondence between how subjects’ behavior suggested they were using the sequences, and the neural representations of the sequences in OFC ensembles. Multiplexed with this value-invariant representation of the task, we also found a representation of the expected value at each location. Thus value and task structure are co-existing and potentially dissociable components of the neural code in OFC.

## INTRODUCTION

It is widely believed that the OFC is part of a neural circuit signaling information about future outcomes (Rudebeck and Murray, 2014). But what is the OFC’s role in that network? What information does it provide exactly? One proposal is that the OFC compresses information about these future events down to their “economic” value (Padoa-Schioppa and Conen, 2017). Another is that the OFC represents a cognitive map of the current state space – a detailed associative model of the causal relationships between events, useful for determining (but not synonymous with) value (Schuck et al., 2016; Wilson et al., 2014). In favor of the former proposal, neural correlates of value are ubiquitous in OFC, generally dominating the neural code in rodents, monkeys and humans (Gottfried et al., 2003; Howard and Kahnt, 2017; Kennerley et al., 2011; Kennerley et al., 2009; Levy and Glimcher, 2011; Padoa-Schioppa and Assad, 2006; Plassmann et al., 2007; Rich and Wallis, 2016; Roesch and Olson, 2004; Roesch et al., 2006; Rudebeck et al., 2013; Thorpe et al., 1983; Tremblay and Schultz, 1999; Zhou et al., 2015). Yet the tasks used in recording studies typically employ designs and heavy training to randomize, trivialize or otherwise make irrelevant everything but the value available on a given trial; when this is not done, neural correlates of value-neutral and even incidental associations are evident (Deikhof et al., 2011; McDannald et al., 2014; Sadacca et al., 2018; Wimmer and Shohamy, 2012). Further, the OFC is generally only necessary for value-based behavior when the relevant value depends on the sort of mental simulation that traditionally is thought to require a cognitive map (Gallagher et al., 1999; Gardner et al., 2017; Izquierdo et al., 2004; Jones et al., 2012; Reber et al., 2017; Schoenbaum et al., 2011).

One way to reconcile these two seemingly opposing ideas is to view them as extreme ends of a continuum determined by the complexity of the environment and the subject’s experience in it. Clearly a naïve subject cannot have much of a cognitive map. However, learning that happens in a few trial (or even a single trial) shows that a cognitive map can be rapidly initialized with potential relevant relationships, and in the initial phase of sensory preconditioning, OFC neurons acquire representations of seemingly incidental sensory-sensory associations (Sadacca et al., 2018). As training proceeds, such a map might be pruned or edited down to increasingly abstract cognitive constructs, according to the subject’s understanding of the structure of the environment at any particular time. The resultant map might appear to represent value in simple situations, while maintaining a great deal of complexity and specificity about prior and future events when these are relevant to the desires of subject in more complex settings.

Here we tested this prediction in rats by recording single unit activity from the OFC during the performance of an odor sequence task (Ginther et al., 2011). The task was based on a standard go/no-go discrimination task (Figure 1A), well documented to engage value coding in the OFC (Critchley and Rolls, 1996a, b; Rolls et al., 1996; Schoenbaum et al., 1998, 1999; Schoenbaum and Eichenbaum, 1995; Schoenbaum et al., 2003). However, rather than presenting the odors pseudorandomly, we presented them in 6-trial sequences, intentionally confounding structural information about sequence and value. There were 4 sequences arranged in two pairs. Each sequence pair was analogous to an inverted T-maze, beginning with a pair of unique “arms” in which the odors differed, followed by a single common “arm” in which the odors were identical (Figure 1B). Conceptually, this resulted in 24 unique “positions” defining the global cognitive map or state space of the task. Within this state space, some positions were defined by external, observable information (unique odors) while others were defined only by reference to internal, unobservable information (memory of the prior sequence of odors). Importantly, in one sequence pair (Figure 1B, top), the shared or common odors were associated with the same actions and rewards, while in the other sequence pair (Figure 1B, bottom), actions and rewards on two of the odors in the middle of the common arm differed depending on which of the unique arms initiated the sequence. As a result of this arrangement, maintaining separate representations of the current position in the sequence was necessary for task performance in some sequences (and positions), whereas in analogous positions in other sequences information about which sequence the position is in was incidental. We analyzed ensemble activity to assess how well OFC represented the unique positions within each sequence and the dependence of any such positional or state representations on external versus internal information and task relevance or value.

**Figure 1.**
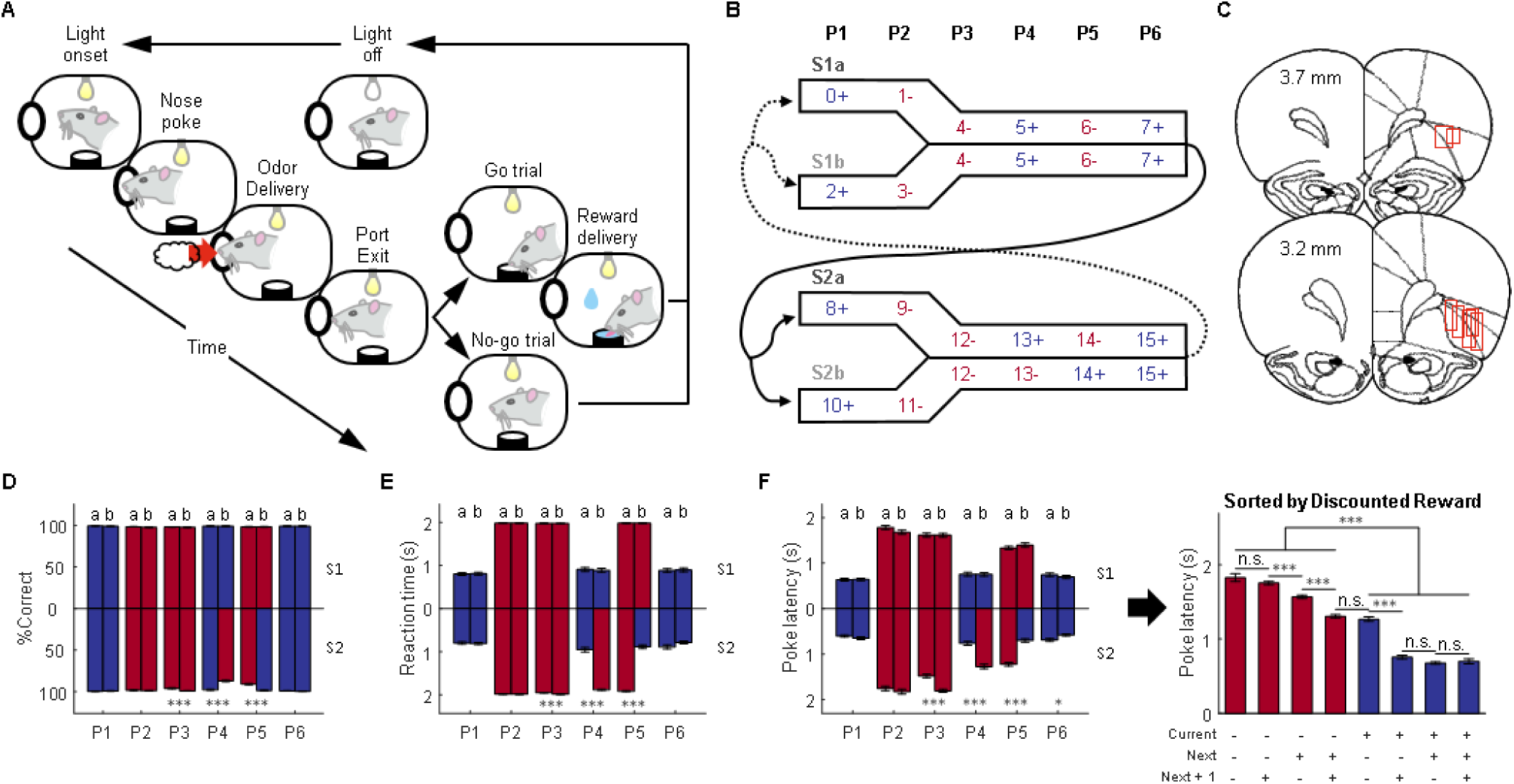
Task design, histology, and behavioral performance. (A) Schematic illustrating a single trial from the odor sequence task. Illumination of an overhead light indicated the start of each trial. After poking into the odor port and sampling one of 16 odors, rats made a “go” or “no-go” decision based on the current odor identity and prior sequence information. Only correct “go” trials led to a liquid reward. (B) The 16 odors were organized into two pairs of sequences (S1 and S2, each comprised of two subsequences a/b). Each sequence consisted of 6 discrete positions marked by the odors (Blue = rewarded, Red = non-rewarded). The odors defining the first two positions (P1, P2) were unique in each sequence, while the odors defining the last four positions (P3-6) were the same across pairs of sequences (S1a/S1b and S2a/S2b). The trial sequence alternated between S1 and S2, with approximately equal transitions between the subsequences. (C) Reconstruction of recording locations in the OFC. Red boxes indicate approximate location of recording sites. We recorded 1,078 single units from 7 rats. (D) Animals’ performance was assessed by percent correct responding (%Correct) on each trial type (n = 73 sessions). At P3, P4, and P5, the percent correct was significantly different between S2a and S2b (p = 1.7 × 10^−5^, 2.2 × 10^−16^, 8.4 × 10^−13^; W = 4472.5, 7386, 3680.5, respectively; two-sided Wilcoxon rank-sum test; n = 73 sessions). (E) Reaction time from withdrawing from the odor port (“unpoke”) to the entry of water port (“choice”). Reaction time on go/reward trials was significantly lower than that on no-go/no reward trials (p = 1.7 × 10^−25^; W = 2701; two-sided Wilcoxon rank sum test). The reaction time was also significantly different between S2a and S2b at P3, P4, and P5 (p = 7.8 × 10^−5^, 1.3 × 10^−20^, 4.2 × 10^−23^; W = 4552, 2996, 7883, respectively; two-sided Wilcoxon rank-sum test; n = 73 sessions). (F) Poke latency from light onset to poking into the odor port measures animals’ motivation to initiate a trial (left panel). The poke latencies were significantly different between S2a and S2b at P3, P4, P5, and P6 (p = 2.6 × 10^−8^, 2.2 × 10^−11^, 4.1 × 10^−15^; 0.013; W = 3942, 3655, 7372, 6001 repectively; two-sided Wilcoxon rank-sum test; n = 73 sessions). Plotting poke latency against time-discounted reward (right panel) showed that poke latency on rewarded trials was significantly less than that on no-reward trials, meaning that rats spent much less time initiating a “go” trial (p = 6.5 × 10^−236^; W = 420567; two-sided Wilcoxon rank sum test; n = 73 sessions). In addition, statistical analyses on adjacent bars showed that poke latency was also negatively modulated by future reward vs. non-reward (p = 0.16, 1.5 × 10^−17^, 5.9 × 10^−15^, 0.64, 6.5 × 10^−27^, 0.065, 0.57; W = 14490, 83436, 68957, 86362.5, 92495, 59118, 52978 for consecutive pairs of bars from left to right, respectively; two-sided Wilcoxon rank-sum test; n = 73 sessions). Further three-way ANOVA analysis revealed that reward on current, next, and next + 1 trials significantly affected poke latency (current: F(1, 1745) = 2.8 × 10^3^, p = 0; next: F(1, 1745) = 128.1, p = 1.0 × 10^−28^; next + 1: F(1, 1745) = 10.1, p = 0.0015). There were also significant interactions between them (current × next: F(1, 1745) = 84.5, p = 1.0 × 10^−19^; current × next + 1: F(1, 1745) = 41, p = 1.9 × 10^−10^; next × next + 1: F(1, 1745) = 15, p = 1.1 × 10^−4^). P1-P6 refers to position in the 6-trial sequences; data from sequence-pair 1 is plotted upward and data from sequence-pair 2 are plotted downward. For panels D-F the error bars are standard errors of the mean across all sessions included in the ensemble analyses (SEMs). *p < 0.05, ***p < 0.001 and Blue = rewarded, Red = non-rewarded.

## RESULTS

### Odor sequence task

Seven rats were trained on the odor sequence task described above, in which knowledge of the position in the sequence was relevant to—and sometimes required for—optimal performance (Figure 1). Rats sampled one of 16 odors on each trial and made a “go” or “no-go” response to obtain reward or to avoid a prolonged inter-trial interval (Figure 1A). The 16 odors were organized into two pairs of 6-trial odor sequences (Figure 1B; sequences S1a/b and S2a/b). S1a and S1b were always followed by S2a or S2b and *vice versa*, and the likelihood of a given transition – such as S1a to S2a versus to S2b - was roughly equal (Figure 1B). The sequences were evenly distributed within each session, and their overall order was the same for all rats in all sessions.

Each sequence pair was intended to function like a maze, with the position being defined by the identity of current and prior odors in each sequence. The odors in the first 2 positions of each sequence (P1 and P2) were unique, like the different arms of a maze, so that they defined a unique position without reference to the identity of the prior odors. By contrast, the odors in the other 4 positions (P3 - P6) were identical in each sequence pair, like the common arm of a maze, so that they defined a sequence-unique position only in concert with the prior odor cues. Critically, in S1a and S1b, these common odors were associated with actions and rewards that depended only on the current odor, whereas in S2a and S2b, opposing actions were required for and different rewards were predicted by two of the common odors (in P4 and P5), depending on previous odors (that is, these odors predicted reward differently in S2a and S2b).

We recorded 1,078 single units from the OFC of rats performing this task (Figure 1C). During recording, rats were generally excellent at the task, responding correctly to the cues at each position in each sequence (Figures 1D and 1E; see Figure S1 for information on pre-recording training and the performance during recording). Behavior was nearly perfect for the odors whose reward predictions did not change across sequences, however the rats also performed well on the odors whose meaning required the rats to use information about the sequences of odors (positions P4 and P5 in bottom bars in Figure 1D). In addition, for all cues, the rats were faster to initiate trials when the sequence predicted reward for that trial than when it predicted no reward (Figure 1F, left panel; note that this is prior to odor onset, so can only reflect past sequence information). This indicates that even when odors uniquely predicted reward, so sequence information was not required for predicting reward, rats were still sensitive to that information and used it to influence their behavior. Indeed, these latencies also showed a significant effect of future reward (Figure 1F, right panel). This general pattern of behavior was exhibited by the group and by each individual rat (see Figure S2 for parallel analyses of sessions from each rat).

### Ensembles in OFC encode both the value and state defining the current trial

We constructed pseudo-ensembles composed of neurons recorded in different sessions and rats and analyzed the ability of the pattern of activity in these populations to correctly identify the position of the current trial within the various sequences. To illustrate the two extreme outcomes of this analysis foreshadowed by our introduction, we plotted the results as they would appear if OFC represented only expected value, defined by the reward available on the current trial, versus the actual map of states in the task, defined by each position in the odor sequences (Figure 2A). In each plot or “confusion matrix”, the Y-axis shows the actual position of a trial in the sequence (P1-P6, and within these, S1a-S1b-S2a-S2b), and the X-axis shows how the ensemble would classify that trial, on average, based on the two encoding schemes. If coding in OFC represents each unique location or state in each sequence, this would result in classification along the diagonal of the matrix (Figure 2A, right), since each position in each sequence is unique due to the odors on the current and/or prior trials. By contrast, if coding is driven by current value, independent of sequence (Figure 2A, left), then trials would frequently classify in other parts of the matrix, reflecting the fact that half the positions are associated with reward. A comparison of these idealized plots with the results of the analyses of firing rate data from different epochs in each trial shows relatively poor correspondence between the raw data and either extreme alternative (Figure 2B; For single-unit examples see Figure S3; For decoding analyses on individual rats see Figure S4). Even during the odor presentation period, there is substantial classification off the diagonal, indicating that raw firing rates are not encoding current location within the sequence with perfect fidelity; on the other hand, even in the actual reward period, there is a strong representation along the diagonal, indicating that raw firing rates are also not encoding reward or current value independent of sequence.

**Figure 2.**
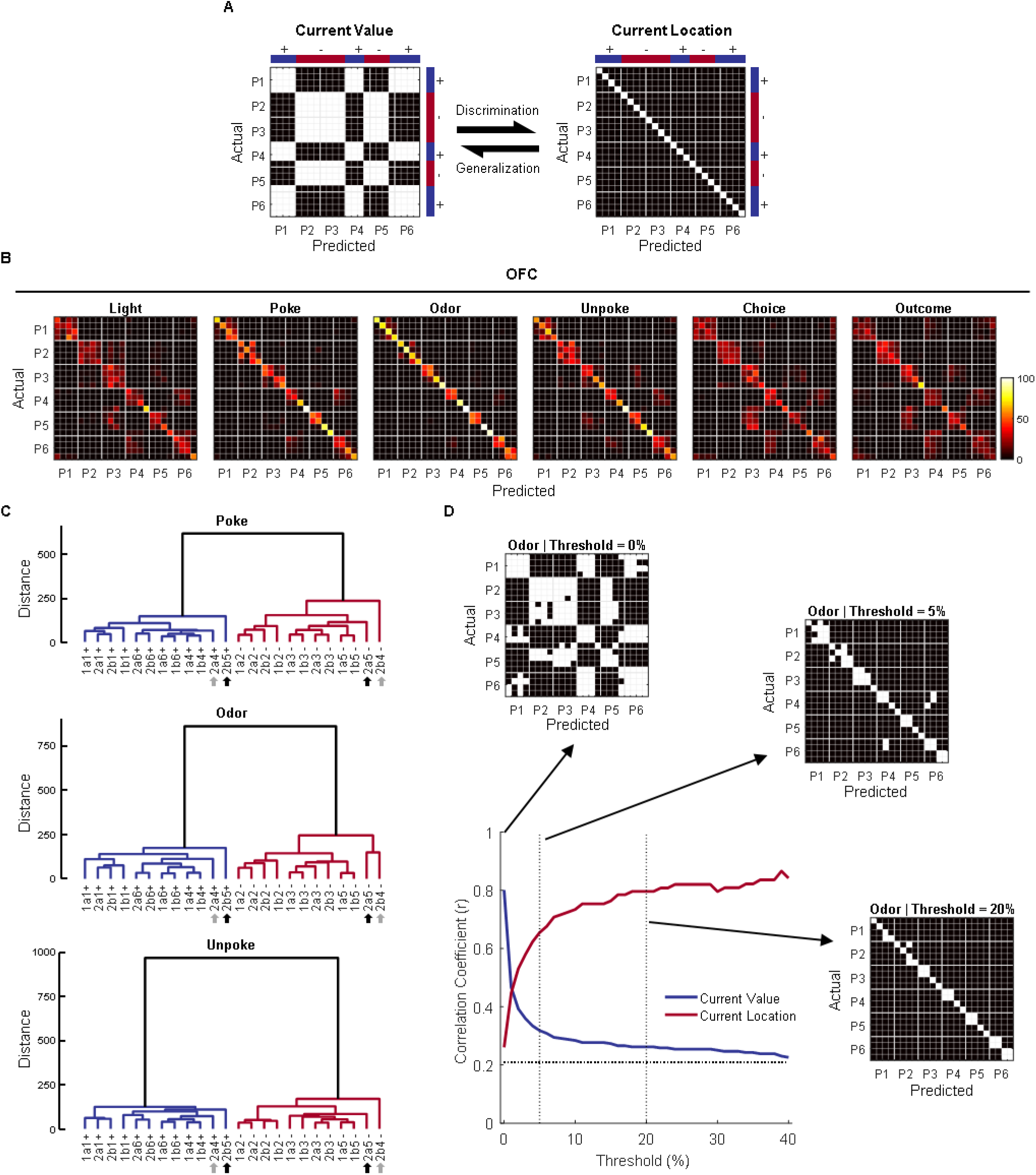
Decoding and clustering of 24 locations. (A) The two confusion matrices represent two hypothesized models (“Current Value” and “Current Location”). The Y-axis represents the ground-truth 24 trial types, ordered by position (P1-P6) within each sequence (S1a, S1b, S2a, S2b). The X-axis represents how these 24 predicted trial types would be classified according to the hypothesized information (e.g. value or location in the sequences). In the left confusion matrix, trial types are misclassified into positions with the same current value, resulting in a checkerboard pattern. By contrast, in the right confusion matrix, each trial type has a unique location or state in each sequence, resulting in perfect classification along the diagonal. (B) Confusion matrices from decoding of 24 trial types from OFC single-unit data in time windows associated with different task events. (C) Dendrograms show hierarchical clustering of 24 locations at 3 different task events (“Poke”, “Odor”, and “Unpoke”) based on population neural activities. The Mahalanobis distance between each pair of trial-type means, which reflects representational dissimilarity or distance between locations, was used to construct the hierarchical clustering tree with an unweighted average linkage method. Shown in the dendrograms, the clustering analyses revealed a detailed relationship between 24 locations represented by the OFC pseudo-ensembles beyond decoding analyses. Grey arrows indicate odor 13 at P4 in S2, and black arrows indicate odor 14 at P5 in S2. Decisions on these odors (4 trial types) require past sequence information. (D) Binarization of confusion matrices using different thresholds. The raw confusion matrix at odor time (shown in B) was filtered at different thresholds (0%, 5%, and 20%; anything above the threshold was painted white and the rest is black) to extract different patterns of information. Value was evident at very low thresholds (0%), whereas detailed location coding was more prominent at higher thresholds (5% and 20%). Line plot shows correlation coefficients comparing the similarity between hypothesized “current value” and “current location” matrices and the actual confusion matrices at different filtering thresholds (0% – 40%). The dotted horizontal line indicates the correlation coefficient between the two extreme models – correlation with either of the models at that level cannot reliably indicate one model rather than the other.

To visualize the underlying structure of the neural activity space that gives rise to the classification patterns in the confusion matrices, we constructed dendrograms summarizing the Mahalanobis distance between each pair of trial types in the ensemble activity space. By plotting these distances in a tree-like structure, we can see how the trial types are clustered and which neural representations are more similar to each other, rather than just the “best match” revealed by the classification in the confusion matrices. The results, shown for the poke, odor, and unpoke periods (Figure 2C), show that current trial value was a major determinant of how the trial types clustered, and therefore how the neurons coded the different trial types. This was evident in the high degree of dissimilarity between the rewarded trial types, in blue, and the non-rewarded trial types, in red, in each dendrogram. However, beneath that global structure, the sequential structure of the state space is well represented. In each period, odors at similar positions in the sequences (P1, P2, P3…) tended to cluster together, indicating that sequence position influenced the representation in the neural activity space. The exception to this organization were the two odors with sequence-unique reward predictions, at P4 and P5 in S2, indicated by the gray and black arrows under the dendrograms, which seem to be represented differently from the other odors (and therefore separate at a higher level in the hierarchy in the dendrograms). Importantly, these value and state features characterized the activity space during, after, and also *before* odor presentation. Representation of information about the trial before odor presentation is consistent with behavioral evidence that the rats used sequence information to anticipate the upcoming trial, even when this was not necessary (Figure 1E-F).

The different levels of information available in the neural activity space, evident in the dendrograms, can also be revealed by filtering the results of the confusion matrices at different thresholds. In this analysis, we set a threshold, say, 5% confusion, and painted as similar (white) states that were confusable at that threshold or above (that is, states that would be classified as identical in more than 5% of cases). A simple value pattern dominated at very low thresholds (Figure 2D, 0%) indicating that if we consider as identical any states that are *sometimes* confused with others, states become grouped by whether they are rewarded or not. Conversely, a pattern more consistent with sequence dominated at higher thresholds (Figure 2D, 5% and 20%), suggesting that the more fine-grained differences between neural representations of different states were able to separate states according to their sequence and location, even when odors were identical, and despite the reward value of these states being similar. A formal analysis tracking the correlation coefficient between the filtered patterns and the two iconic exemplars (Figure 2A) at different levels of filtering confirmed this impression, showing that information about value was available only at the lowest filtering levels, declining precipitously even at a threshold of 2-3%, whereas information about structure increased quickly, overtaking value and remaining high through a large range of filtering thresholds (10-40%; Figures 2D and S5).

### Representations of current trial value and state in OFC are dissociable

The analysis presented above shows that the OFC contains information relevant to value but that this information is embedded within a rich representation of task structure. To test whether the neural codes for value and task structure were dissociable, we utilized a linear discriminant analysis (LDA) to isolate different components explaining the variance across the pseudo-ensembles. First, the firing rates of 1078 recorded neurons on each trial, constituting a 1078-dimensional vector (360 trials in total; 24 trial types;15 correct trials for each trial type), were reduced to a 151-dimensional space though principle component analysis (PCA; the first 151 principal components explained 80% variance). The LDA analysis then transformed these principal components to an equal number of orthogonal LDA components (Figure 3A), ordered by how much of the reward variance they explained. The resultant first LDA component perfectly separated the trial types based on current trial value (Figure 3B), while the other 150 components exhibited no selectivity for current value at either the level of the individual components (Figure 3C) or in the aggregate (Figure 3D).

**Figure 3.**
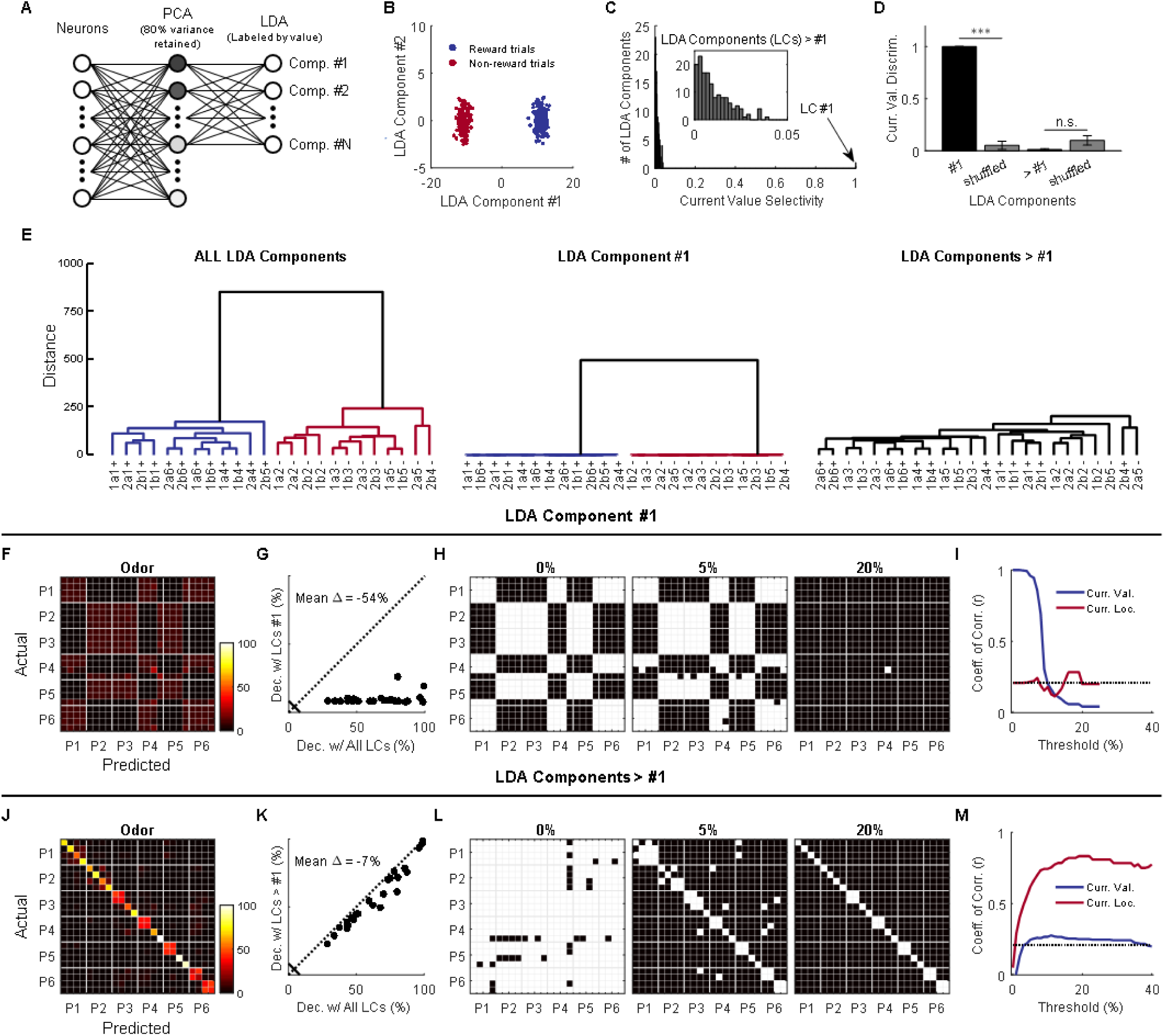
Value and state representations are dissociable at the neural ensemble level. (A) The firing rates of all single neurons on each trial constituted a high-dimensional vector (360 vectors or data points in a 1078-dimentional space). The firing rates of all neurons at the odor time were linearly projected to a principal component subspace with 80% variance explained, then to an LDA space with labels about the current reward. Each LDA component combined a weighted sum of inputs from all the neurons. The LDA transformation was supervised by trial-type labels that only separated current value (reward vs. non-reward), so that the LDA could find components that best separated the two classes. Comp., Component. (B) The first but not the second LDA component perfectly separated the two trial types (p = 1.0 × 10^−3^ and 1.0, respectively; two-sided permutation test, 1000 bootstrap samples). (C) An ROC-based value-selectivity index (2 × |AUC – 0.5|) ranging from 0 (low selectivity) to 1 (high selectivity) was used to test current value selectivity for each individual LDA component. The first LDA component showed perfect value selectivity (1.0; p = 1.0 × 10^−3^; permutation test; 1000 bootstrap samples). But, none of the remaining 150 LDA components were selective for the current value (< 0.05; p = 1.0 for all components; two-sided permutation test; 1000 bootstrap samples). (D) Value discriminability was used to test whether value was distributed across components (0 – 1 indicates the level of value discriminability by population components). The true discriminability was compared with that from the label-shuffled data. The first LDA component showed significantly higher value discriminability than the shuffled data (1.0 vs. 0.05; p = 1.0 × 10^−3^; one-sided permutation test; 1000 bootstrap samples), but the remaining LDA components did not show significant higher value discriminability than the shuffled data (0.13 vs. 0.1; p = 1.0; one-sided permutation test; 1000 bootstrap samples). Curr. Val. Discrim., Current Value Discriminability. Error bars are standard deviations (SDs). (E) A Dendrogram using all different LDA components contained both value and state information (left). A dendrogram that only used the first LDA component only contained value information without detailed state information (center), while a dendrogram that only used the remaining LDA components contained state information without current value (right). (F) Decoding of 24 states with the first LDA component (reconstructed to 151 PCs before the decoding analysis). (G) Comparison of decoding accuracy for each state (represented by each dot) between all LDA components and the first LDA component being used. Dec. Decoding; LCs, LDA Components. (H) Confusion matrix at odor time was binarized at thresholds 0%, 5%, and 20%. (I) Correlation coefficients compare the similarity between hypothesized “current value” and “current location” matrices and the actual confusion matrices (obtained by using the first LDA component) at different filtering thresholds. (J) Decoding of 24 states with the remaining 150 LDA components (reconstructed to 151 PCs). (K) Comparison of decoding accuracy for each state between all LDA components and the remaining LDA components (the first one was left out) being used. (L) Confusion matrix at odor time was binarized at thresholds 0%, 5%, and 20%. (M) Correlation coefficients compare the similarity between hypothesized “current value” and “current location” matrices and the actual confusion matrices (obtained by using the remaining LDA components) at different filtering thresholds.

This suggests that the representation of value in the population could be orthogonal to the representation of the sequence structure – not at the level of individual single units but rather in the overall pattern of their firing – and therefore the two representations are effectively multiplexed in the neural signal. Such orthogonalized representations may be utilized downstream, allowing different brain areas to easily access different aspects of the information output by OFC. To assess this, we repeated the analyses done on the raw data in Figure 2, isolating firing rates derived from the first LDA component versus the remaining value-neutral LDA components. These analyses cleanly dissociated the value and structure encoding that was confounded in the analysis of the raw data. Thus a dendrogram produced using neural data transformed by the first LDA component retained the separation based on current trial value but lost nearly all of the underlying structure (Figure 3E, left vs. center), whereas a dendrogram produced using neural data transformed by the remaining components retained the sequence structure but lost all information reflecting current trial value (Figure 3E, left vs. right). Further, the confusion matrices produced by the two transformed data sets hewed closely to the iconic patterns at the top of Figure 2; filtering of these patterns revealed value encoding in analyses of the first LDA component with no information about structure (Figure 3F-I), and structure encoding in analysis of the remaining components with no information about value (Figure 3J-M).

### Ensembles in OFC encode sequence information across trials when task relevant

In addition to representing current position in the sequence, the OFC also maintained sequence information across positions, depending on task relevance. To show this, we trained a classifier using neural activity excluding the first LDA component at each position (P1 – P6), and then used it to decode activity from trials at other positions (P1 – P6) (Stokes et al., 2013). The results of this cross-positional decoding are illustrated in matrices for sequence S1 (Figure 4A; center) and S2 (Figure 4B; center). The diagonals of the two matrices show how well each position can be decoded using training data from itself (Figure 4A and B; upper-left). These plots confirm the results of the earlier analyses, showing that activity in OFC is able to distinguish positions well when prompted by either external information (P1 or P2 in both sequences) or task relevance (P3, P4, P5 in sequence 2). However, the off-diagonal cross-positional decoding also shows that the representation of information about sequence extended across trials in the sequence. That is, for some positions, the classifier built upon the data from one position in the sequence could be used to correctly decode data from other positions earlier or later in the same sequence. Cross-positional decoding was particularly prominent when remembering position was necessary for correct performance in S2 at the transition point into the common arm, at P2 – P4 (Figure 4B; center and right). Notably, there was no such cross-positional decoding at the transition point and in the common arm of S1 (Figure 4A; center and right). This difference is particularly evident if one focuses on the two rows in which the classifier was trained with data from P4 or P5 and used to decode sequence at the prior positions (Figures 4A and B, right). This analysis revealed above-chance decoding at earlier positions in S2, whereas not a single position was decoded above chance in S1. Interestingly, the classifier trained at P4 did a poor job decoding at P5 and *vice versa*; but both P4 and P5 showed good cross-positional decoding at P3. This suggests the existence of two orthogonal persistent codes at P3 (shared with P4 and P5, respectively), which together help rats keep track of different outcomes or rules in the sequence.

**Figure 4.**
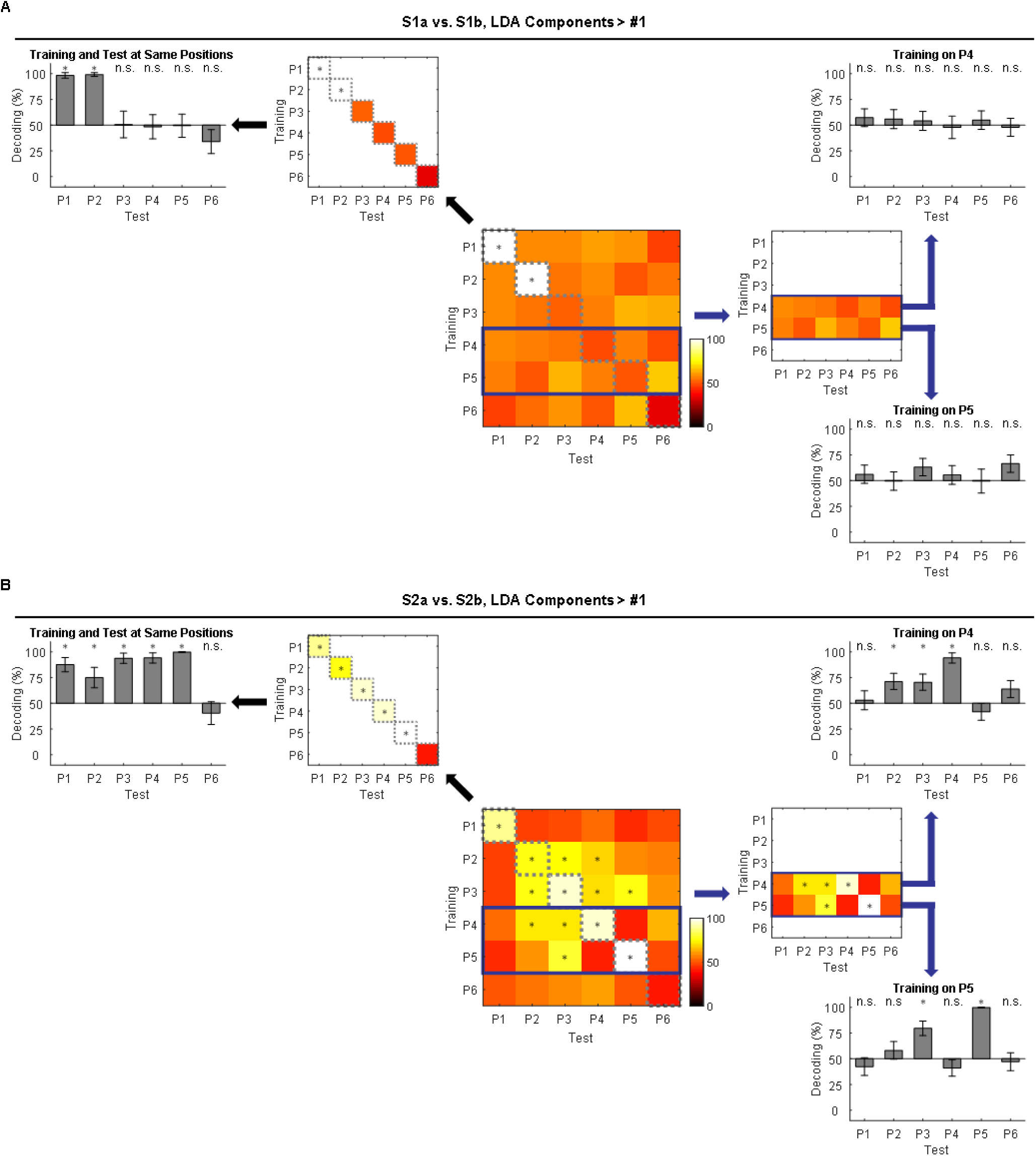
Cross-positional decoding of sequence information. (A) Cross-positional decoding of sequence S1a vs. S1b. A classifier to discriminate S1a vs. S1b at odor time was trained at each position (P1 – P6) and tested at other positions (P1 – P6). Decoding accuracy on the diagonal was reproduced and plotted to its upper-left. A bar graph on the left summarized the data in the diagonal and showed how well S1a vs. S1b was decoded at each position (P1 – P6; statistical significance was determined by the mean decoding accuracy being outside the 95% CIs estimated by the same decoding process with label-shuffled data). On the right side, two rows (classifiers trained at P4 or P5, but tested at P1 – P6) were highlighted in the heatmap and summarized as bar graphs (P1 – P6; significance was determined by 95% CIs). (B) Cross-positional decoding of 2a vs. 2b. The data are displayed in the same format as in (A). Error bars are SDs.

### Ensembles in OFC miscode sequence when the rat miscodes sequence

The analyses to this point suggest that neural activity in the OFC is shaped by the task. Although current trial value is a major determinant of this activity pattern, when information about the trial structure or sequence is important for correct performance, this information is also maintained. If this is true, then one might expect activity on error trials – when the rats make a mistake in deciding whether to go or not go – to reveal miscoding in the activity space. In our task, mistakes were almost completely restricted to P4 and P5 in S2, which required rats to recall the sequence of prior trials to respond correctly (Figure 1D). Errors of commission on P4 trials are particularly useful for this analysis, since unlike P5, there was no information available for several prior trials regarding whether the rat was in S2a or S2b. Thus it is possible to ask whether sequence was miscoded on the error trial as well as on the preceding trial, without any interference from outside input. Further, the nosepoke latencies at P4 in S2 suggested that mistakes at P4 typically occurred because the rat believed it was in one sequence when it was actually in the other. Thus the rat would initiate an error trial in one sequence with a latency appropriate for the correct trial in the other (Figure 5A).

**Figure 5.**
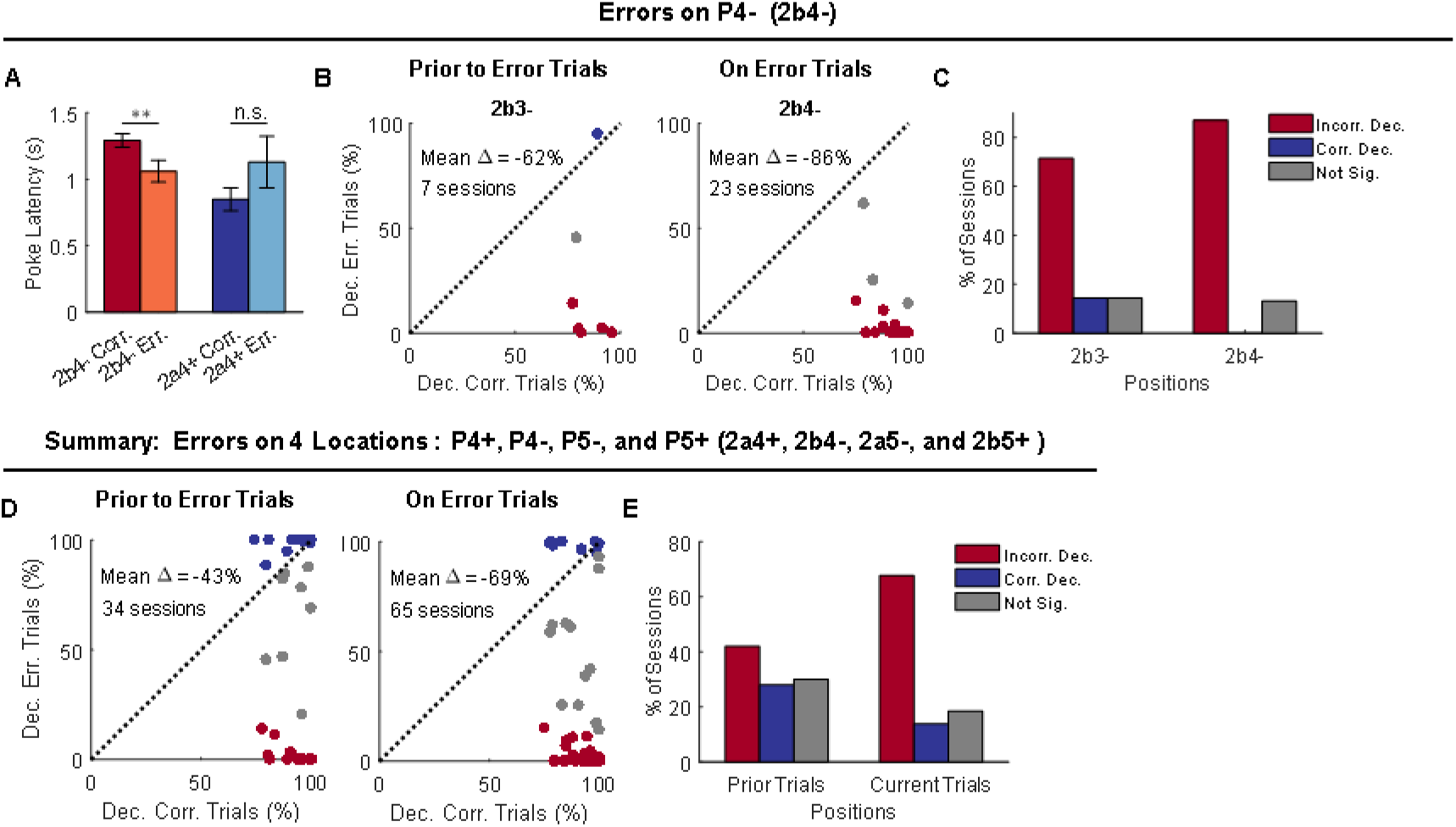
Decoding of sequences at current and prior positions on current correct versus error trials. (A) Poke latencies on 2b4-correct and error trials were significantly different (p = 0.003; W = 1524; paired, two-sided Wilcoxon rank sum test; n = 65 sessions), but those on 2a4+ correct and error trials were not significantly different (p = 0.15; W = 100; paired, two-sided Wilcoxon rank sum test; n = 24 sessions). Two-way ANOVA analysis revealed an interaction effect between sequence (2a4+ vs. 2b4-) and performance (correct vs. error; F(1, 174) = 6.37, p = 0.013). Error bars are SEMs. (B) Classifiers were trained with only correct trials and tested with correct trials (X-axis) or error trials (Y-axis). The decoding analyses were carried out on both current error trials (2b4-) and on trials that prior to error trials (2b3-). Only sessions that had error trials on 2b4- and also showed above-chance decoding of 2a vs. 2b at P3 or P4 (mean chance level: 50%; significance was determined by 95% CIs) were selected for further analyses. The total number of sessions is indicated in plots. Blue dots indicate significantly above-chance decoding of 2a vs. 2b for error trials (mean chance level: 50%; significance was determined by the right side of 95% CIs). Red dots indicate significantly below-chance decoding of 2a vs. 2b for error trials (mean chance level: 50%; significance was determined by the left side of 5% CIs). Gray indicate non-significance (within 95% CIs). (C) Percentage of sessions that show significantly below- (red), above- (blue), or non- (gray) significant decoding of 2a vs. 2b prior to error trials (dependence between behavioral and decoding performance: χ^2^ = 10.5; p = 0.0052; chi-squared test) and on error trials (χ^2^ = 46.0; p = 1.0 × 10^−10^; chi-squared test). (D) Summary of decoding analyses on four error trials (2a4+, 2b4-, 2a5-, and 2b5+). Decoding of 2a vs. 2b at prior position with correct (X-axis) or error trials (Y-axis) in the current trials (left panel). Decoding of 2a vs. 2b at current position with correct (X-axis) or error trials (Y-axis) in the current trials (right panel). (E) Percentage of sessions that show significantly below-(red), above-(blue), or non-(gray) significant decoding of 2a vs. 2b prior to error trials (χ^2^ = 56.3; p = 6.1 × 10^−13^; chi-squared test) and on error trials (χ^2^ = 98.4; p = 4.3 × 10^−22^; chi-squared test). Incorr. Dec., Incorrect Decoding; Corr. Dec., Correct Decoding; Not Sig., Not Significance.

These error trials and their behavioral similarity to correct trials on the opposite sequence in a pair give us a unique opportunity to ask whether encoding of information about sequence in the OFC is mere happenstance or if it is directly related to what information the rat is acting on versus what it is experiencing – that is, does it reflect the hidden variable of state? To test this, we compared how well ensembles recorded in individual sessions performed at decoding sequence on correct versus error trials in S2 (Figure 5). We focused on decoding on correct and error trials at S2b4-, which is best positioned to address this question, and restricted the analyses to individual sessions in which simultaneously recorded ensembles exhibited above-chance decoding on correct trials (Figure 5B and C). Results showed that while these OFC ensembles represented a trial as belonging to the S2b4-when the rat responded correctly, they represented the trial as belonging to the opposite sequence (S2a4+) when the rat responded incorrectly. Further the miscoding was present both on the actual error trial (P4; S2a4+ vs. S2b4-) and also on the trial preceding the error trial (P3; S2a3+ vs. S2b3-). Similar miscoding was also observed on a substantial number of such runs of trials involving errors at the other positions (Figure 5D and E).

## DISCUSSION

Historically, the OFC has been implicated in signaling information about expected outcomes relevant to ongoing adaptive behavior (Jones and Mishkin, 1972; Rolls, 1996). Current proposals contrast signaling of expected value with representing a cognitive map of the task (Padoa-Schioppa and Conen, 2017; Wilson et al., 2014). However, these two proposals are not mutually exclusive. Beliefs regarding the associative structure of a task are critical to determining the value of the current trial, while the value of the current trial is a critical component of the underlying task structure. In a well-trained subject, the externally available information should reflect the abstract task-relevant cognitive constructs – underlying hidden states – formed with experience. This would result in a cognitive map suitable to the subject’s decision-making needs in a given task. That is, states of a task should be compressed or represented separately in OFC based on whether distinguishing them is important for current behavior. In simple tasks, in which individual trials and their outcomes are isolated, like those generally used for single-unit recording, the final product might appear to only represent value, but in more complex situations in which choices are made in the context of ongoing behavior, the representation should maintain a complexity that matches the behavioral strategy of the subject and the causal relationships in the task.

Here we tested this prediction by recording single unit activity from the OFC in rats performing an odor sequence task that provided a complex but mappable state space spanning sequences of trials. These sequences could be thought of as analogous to a spatial maze, with individual trial types reflecting distinct locations in the maze. Consistent with the above hypothesis, we found a close correspondence between how the subjects’ behavior suggested they were mapping the sequences and the neural representations of the sequences in OFC ensembles. Specifically, neural ensembles distinguished positions in the sequences in the complete absence of any externally distinguishing information when such discrimination was necessary for the behavior of the rat on the current or subsequent trials (S2 common arm); similar positions were not distinguished when there was no behavioral relevance to their distinction (S1 common arm). This was true for the current position in the sequence and also for decoding of other nearby positions, and the representations were faithful to the rats’ internal classification of which sequence they were in, such that when the rats’ behavior seemed to miscode the sequence, the ensembles in OFC miscoded the sequence as well. Interestingly, the ensembles also distinguished positions when external information was sufficient to do so (unique arms), and even when doing so was seemingly irrelevant to the rats’ current or future behavior (S1, unique arms). This suggests that some external information may be too salient to fully compress and ignore, or that the rats are using this information in ways we cannot appreciate with our response measures. However, the general pattern of neural activity was largely consistent with the idea that OFC represents the cognitive map of the trial structure necessary for the behavior or actions of the subject.

Our results also show that information about current position in the sequence could be formally dissociated from information about current value, not in the raw data but in a linearly-transformed neural activity space. This analysis reduced a ∼1000-neuron population to a much smaller number of components, and within these, a single component encompassed the bit of information relevant to the value of the current trial, while the remaining components contained the much more detailed information about the sequence of trials in which the value was embedded. While value is often found to co-occur with encoding of other information in OFC at the level of single units or populations (Howard and Kahnt, 2017; Kennerley et al., 2011; Kennerley et al., 2009; Padoa-Schioppa and Assad, 2006; Roesch et al., 2006; Rudebeck et al., 2013; Thorpe et al., 1983; Tremblay and Schultz, 1999), this is rarely highlighted (but see Farovik et al., 2015; and Yang and Murray, 2018). Showing this co-occurance clearly and in a complex setting, and showing that the two codes are dissociable has important implications. First and foremost, this result shows how overwhelming value information is, even in the context of an informationally complex task. Value accounted for the largest amount of variance across our population of neurons. In a simpler task, without the trial-spanning structure in our design, value might appear to be the only information of any importance at the ensemble level. However, as value becomes increasingly embedded within a complex associative structure, it may become a smaller component of the activity in the OFC. Second, the representation of task or associative structure was dissociable from value. It is not secondary to or dependent on value; rather it exists in OFC even when value is formally irrelevant or entirely absent (McDannald et al., 2014; Sadacca et al., 2018; Schuck et al., 2016). If the information is dissociable in principle, by such a simple linear analysis, then downstream brain regions could also dissociate these multiplexed signals. Thus output from the OFC could be important not only for providing value predictions to some downstream operator but also for providing a more detailed accounting of the reason why that value was assigned – that is, passing on a picture of the activity space invariant to the distortion of value – to other regions. In this regard, the potential dissociation of these signals provides a simple solution to reconcile the dichotomous views of this area.

## EXPERIMENTAL PROCEDURES

### Subjects

Male Long-Evans rats (Charles River, 175 – 200 g, ∼3-month-old) were housed individually on a 12-h light/dark cycle with ad libidum access to food in an animal facility that was accredited by the Association for Assessment and Accreditation of Laboratory Animal Care (AAALAC). Water restriction was used to motivate rats to perform the task. After training or recording sessions, each rat received 10 min free access to water in their home cages. All testing was conducted at the NIDA-IRP. Animal care and experimental procedures complied with US National Institutes of Health (NIH) guidelines and were approved by National Institutes on Drug Abuse Intramural Research Program (NIDA-IRP) Animal Care and Use Committee (ACUC).

### Surgery

Rats were implanted with drivable bundle of 16 nickel-chromium wires (25 µm in diameter; AM Systems) that targeted the left lateral OFC (AP: 3 mm. ML: 3.2 mm). Wire bundles were housed in a thin cannula and cut with surgical scissors to extend 1.5 – 2 mm beyond the cannula. The tips of wires were initially placed at 4 mm ventral from brain surface, and then driven down 40 µm or 80 µm after each recording session to search for new units. After surgery, rats were give Cephalexin (15 mg/kg po qd) for two weeks to prevent any infection. At the end of testing, rats were euthanized by overdose of isoflurane. The final positions of electrodes were marked by passing a small constant current through the wires, and the brains were processed for histological examination using standard techniques.

### Behavioral testing

Rats were placed in aluminum chambers (∼18” on a side), which were equipped with an odor port and a well for fluid delivery. Behavior was controlled by custom software written in C++ that could monitor responses at the port and well via infrared beam sensors and deliver odors and water by gating a custom-designed system of solenoids. Trial availability was signaled by the illumination of paired house lights above the odor panel, after which the rat had 5 s to initiate a trial by nosepoking at the odor port. If a nosepoke was detected then, after a 500 ms delay, odor was delivered to the port as long as their noses were in the odor port. If the rat left the port in less than 500 ms, the trial was aborted, and the house lights were extinguished. Otherwise, at the end of the odor delivery, the rat had a 2-second time window to respond at the fluid well. On rewarded trials, responding at the fluid well led to delivery of 50 µL of 10% sucrose solution after a delay of 400 – 1500 ms. After the rat consumed the reward and left the well, the house lights were extinguished to end the trial, beginning the ITI. If the rat failed to respond in the 2 s window, the house lights were extinguished at the end of the 2 s period. If the rat responded in the 2 s window on a non-rewarded trial, an exceptionally rare event in recording, the house lights were extinguished, ending the trial, and no reward was delivered. The ITI was 4 s following correct go or no-go trials, and 8 s following trials on which the rat made an error.

One of 16 odors was presented on each trial, and the trials were organized into two pairs of sequences (S1a, S1b, S2a, and S2b). Odors used in each sequence and their associated valence is listed as below.

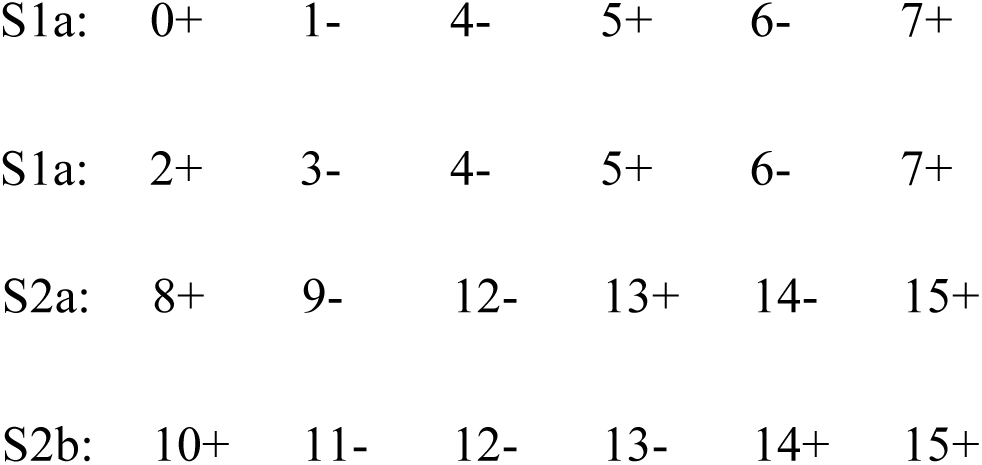

Before training on the full sequence task, rats were first shaped to nosepoke at the odor port and then respond at the well for reward. After this, they were trained to discriminate a single odor pair (one rewarded and one non-rewarded odor) from sequence 1a or 1b. Sessions consisted of a maximum of 480 trials. After rats reached a high criterion in performance (> 90% correct ratio), additional odor pairs were added until the rats were able to perform well in a session containing sequences 1a and 1b. After learning sequences 1a and1b, rats were trained to discriminate between odors 13/14 from sequence 2, including several reversals of the valence of the pair. After the third reversal, additional odor pairs were added from sequence 2 if the rats were able to maintain accurate performance (> 75% correct) on each trial type. Once sequence 2 had been fully introduced in this manner, the rats began sessions containing both pairs of sequences (1a, 1b, 2a, and 2b).

In this final phase, each sequence (or each of the 24 trial types) was repeated for 20 times to make up 480 trials in total. Sequences 1a and 1b were always followed by 2a or 2b with roughly equal probability (0.55 and 0.45, respectively). Sequence 2a and 2b were always followed by 1a or 1b also with slightly more dissymmetry in probability (0.67 and 0.37, respectively). The overall sequence was repeated from start to finish in each session.

|  |  |  |  |  |  |  |  |  |  |  |  |  |
| --- | --- | --- | --- | --- | --- | --- | --- | --- | --- | --- | --- | --- |
| 1b | 2a | 1a | 2a | 1a | 2b | 1b | 2b | 1b | 2b | 1b | 2a | 1b |
| 2b | 1a | 2a | 1a | 2b | 1a | 2a | 1a | 2b | 1b | 2a | 1b | 2a |
| 1b | 2b | 1b | 2a | 1b | 2b | 1a | 2a | 1a | 2b | 1a | 2a | 1a |
| 2b | 1a | 2a | 1b | 2a | 1b | 2a | 1a | 2b | 1a | 2a | 1a | 2a |
| 1a | 2b | 1b | 2b | 1b | 2b | 1b | 2b | 1b | 2a | 1a | 2a | 1a |
| 2b | 1b | 2b | 1b | 2b | 1b | 2a | 1b | 2b | 1a | 2a | 1a | 2b |
| 1a | 2a |  |  |  |  |  |  |  |  |  |  |  |

Before electrode implantation, animals were kept training on the full sequence for at least three weeks until they were able to perform well (> 75% correct) on every trial type.

### Single-unit recording and analyses

Spiking activity was recorded using the Plexon Multichannel Acquisition Processor (MAP) system (Plexon, Dallas, TX). Analog signals from electrodes were amplified (headstage: 20×, differential preamplifier: 50×, acquisition processor: 1 – 32×) and filtered (250 – 8, 000 Hz) following standard procedures. A pre-set threshold was used for each active channel to capture unsorted spikes. Timestamps for behavioral events from the behavioral program were sent to the Plexon system, synchronized and recorded alongside the neural activity. Spikes were sorted to identify single units offline using Offline Sorter (Plexon, Dallas, TX) with a template matching algorithm. Sorted files were opened in NeuroExplorer (Nex Technologies, Colorado Springs, CO) to extract unit and behavioral event timestamps, which were then exported as MATLAB (MathWorks, Natick, MA) formatted files for further analysis. Sample sizes (number of rats and number of neurons) were not predetermined by any statistical methods but are comparable to those reported in previous publications from our lab. All data were analyzed using MATLAB (MathWorks, Natick, MA).

Task events for each trial were synchronized and recorded alongside the neural signals. For neural analyses, each trial was segmented into 6 epochs associated with different trial events: “light”, “poke”, “odor”, “unpoke”, “choice”, and “outcome”. On trials where the rat did not respond, and/or the water reward was not delivered, we used the end of the 2-second window for responding as the “choice” event and a time point 1.5s after the “choice” as the “outcome”. Behavioral performance was quantified as the percent of trials on which the rats responded correctly, their reaction time from the odor port to the fluid well, and the latency with which they initiated a trial after light onset. The data analysis only included sessions in which the percent correct was above 75% for each individual trial type. Error trials were removed in the main figures except Figure 5 in which error trials were used as test sets for classification. The spike train for each isolated single unit was aligned to the onset of each task event. Pre-event time was set to be 200 ms, and post-event time was set to be 600 ms. Spike number was counted with a bin = 100 ms. A gaussian kernel (σ = 50 ms) was used to smooth the spike train on each trial.

### Classification analyses

We trained a linear discriminant analysis (LDA) algorithm (MATLAB function: *fitcdiscr*) to classify 24 trial types or locations for each one of six task events. Neurons that were recorded from different sessions were aligned together as pseudo-ensembles. Firing rates on each trial 100 – 600 ms after task events (500 ms) for each individual neuron were used for classification. Each trial was an observation that contained firing rates from 1, 078 neurons (480 trials in total with error trials; 360 trials in total without error trials; only correct trials were used to build classifiers). The classification accuracy was assessed by leave-one-out cross-validation. Specifically, one trial from each trial type was left out for future testing, and all other trials were used for the training. Principal component analysis (PCA) was used for feature extraction and dimension reduction in the training set. The classifier was trained on the first a few principal components (PCs) that explained 80% variance. The same PCA transformation from the training set was applied to the test set. Trial order for each neuron was shuffled to remove the temporal structure and correlation between neurons within the same trial type. The trial-order shuffling was repeated for 100 times. For each time of trial-order shuffling, the leave-one-out cross-validation was repeated for 500 times. The mean decoding accuracy for each trial type as shown in the confusion matrix was the mean from all runs. The statistical significance of the mean decoding accuracy was determined by 95% confidence interval estimated by running the same decoding process with label-shuffled data. Because of trial order shuffling, the number of PCs being used in classifiers varied for each run. The mean number of PCs being used for each run was 151.2 ± 0.4 (mean ± SD). It was for simplicity when we said 151 PCs were used. To better visualize and potentially extract different aspects of information from the confusion matrices, we binarized the confusion matrices with different thresholds (0% – 40%). For each threshold (e.g. 5%), any value in a confusion matrix that was below or equal to this threshold was set to be 0%, and other places were set to be 100%.

For the classification of all trial types with different LDA components (the first LDA component or the remaining LDA components), we first did PCA on all the trials and neurons (without splitting the data into training sets and test sets) with 80% variance retained (151 PCs). The LDA was run on these PCs, supervised by the value of the current trial (reward vs. non-reward). Thus, each LDA component was a linear combination of original firing rates from all the neurons. Importantly, only the first LDA component showed the ability to discriminate the current value. We reconstructed two sets of PCs from the first LDA component and the other LDA components, respectively. We then used the two sets of PCs to decode the current value (reward vs. non-reward) and 24 trial types (locations or states) with a leave-one-out cross-validation procedure as described above (without further PCA for dimension reduction).

We removed the first LDA components and used others (150 components) for the cross-positional decoding. PCA was used for dimension reduction (with 80% variance retained). We trained binary classifiers at one position (P1 – P5; S1a vs. S1b or S2a vs. S2b) as described above, and tested them on other positions (P1 – P5). The trial orders for both the training and test sets were shuffled. Leve-one-out cross-validation was used for the estimation of the mean decoding accuracy (500 repeats). For each repeat, trials that were left-out for the test set would not be in the training set. Statistical significance was determined by 95% confidence interval estimated from label-shuffled decoding processes.

For the decoding of sequences on error trials in each session, we built classifiers with 15 correct trials and used the error trials as the test set. The trial order was shuffled within each trial type for each repeat. Cross-validation followed the above procedure. The 95% confidence interval was estimated by decoding of sequences on correct trials with shuffled labels in the training set. The mean decoding accuracy was the mean across 500 runs. Chi-squared test was used to test whether there was dependence between behavioral (correct or error trials) and decoding performance (significantly above-chance, below-chance, or not significant).

### Hierarchical clustering analyses

The hierarchical agglomerative clustering was performed on data that was projected onto the LDA space. Each trial was organized as a vector with firing rates of 1, 078 neurons as 1, 078 dimensions. The original data was transformed to PCA space with 80% variance retained (151 PCs), then transformed to LDA space guided by 24 trial-type labels. A dissimilarity matrix was computed by measuring the Mahalanobis distances between each pair of location means. Based on the dissimilarity matrix, an agglomerative hierarchical cluster tree was generated with the unweighted average distance method.

### Discriminability analyses

We used a ROC-based metric to measure how well neural activity components, that were transformed from single unit, can discriminate two different value conditions (i.e. to test whether discrimination of value was distributed across components). The population activity was organized as a matrix of component activity with each row represented one trial (observation) and each column represented a component (dimension). Each trial was labeled with “reward” or “non-reward”. The first three PCs were used for the LDA transformation. We projected all the data points (trials) with value labels onto the first LDA dimension, which was supposed to be the best component to separate value. An ROC curve was constructed to compare the distributions of the neural responses to the two value conditions on the chosen LDA dimension. We used the area under the ROC curve (AUC) to compute the final discriminability metric: 2 × |AUC-0.5|. In addition, we used the same procedure to calculate discriminability of the ensembles on value with shuffled labels, which gave us an estimated baseline discriminability. One-sided permutation test with 1000 bootstrap samples was used to test statistical significance in mean difference between the actual and baseline discriminability (*p < 0.05).

## AUTHOR CONTRIBUTIONS

JZ and GS designed the experiments, within advice and input from SJR and YN. JZ collected and analyzed the data, with advice and assistance from MPHG, TAS, and AMW. JZ and GS wrote the manuscript with input from the other authors.

## ACKNOWLEDGEMENTS

The authors would like to thank Hannah Batchelor for her help with the histology. This work was supported by the Intramural Research Program at the National Institute on Drug Abuse. The opinions expressed in this article are the authors’ own and do not reflect the view of the NIH/DHHS. The authors have no conflicts of interest to report.

## REFERENCES

Critchley, H.D., and Rolls, E.T. (1996a). Hunger and satiety modify the responses of olfactory and visual neurons in the primate orbitofrontal cortex. Journal of Neurophysiology 75, 1673–1686.

Critchley, H.D., and Rolls, E.T. (1996b). Olfactory neuronal responses in the primate orbitofrontal cortex: analysis in an olfactory discrimination task. Journal of Neurophysiology 75, 1659–1672.

Deikhof, E.K., Falkai, P., and Gruber, O. (2011). The orbitofrontal cortex and its role in the assignment of behavioural significance. Neuropsychologia 49, 984–991.

Farovik, A., Place, R.J., McKenzie, S., Porter, B., Munro, C.E., and Eichenbaum, H. (2015). Orbitofrontal cortex encodes memories within value-based schemas and represents contexts that guide memory retrieval. Journal of Neuroscience 35, 8333–8344.

Gallagher, M., McMahan, R.W., and Schoenbaum, G. (1999). Orbitofrontal cortex and representation of incentive value in associative learning. Journal of Neuroscience 19, 6610–6614.

Gardner, M.P.H., Conroy, J.S., Shaham, M.H., Styer, C.V., and Schoenbaum, G. (2017). Lateral orbitofrontal inactivation dissociates devaluation-sensitive behavior and economic choice. Neuron 96, 1192–1203.

Ginther, M.R., Walsh, D.F., and Ramus, S.J. (2011). Hippocampal neurons encode different episodes in an overlapping sequence of odors task. Journal of Neuroscience 31, 2706–2711.

Gottfried, J.A., O’Doherty, J., and Dolan, R.J. (2003). Encoding predictive reward value in human amygdala and orbitofrontal cortex. Science 301, 1104–1107.

Howard, J.D., and Kahnt, T. (2017). Identity-specific reward representations in orbitofrontal cortex are modulated by selective devaluation. Journal of Neuroscience 37, 2627–2638.

Izquierdo, A.D., Suda, R.K., and Murray, E.A. (2004). Bilateral orbital prefrontal cortex lesions in rhesus monkeys disrupt choices guided by both reward value and reward contingency. Journal of Neuroscience 24, 7540–7548.

Jones, B., and Mishkin, M. (1972). Limbic lesions and the problem of stimulus-reinforcement associations. Experimental Neurology 36, 362–377.

Jones, J.L., Esber, G.R., McDannald, M.A., Gruber, A.J., Hernandez, G., Mirenzi, A., and Schoenbaum, G. (2012). Orbitofrontal cortex supports behavior and learning using inferred but not cached values. Science 338, 953–956.

Kennerley, S.W., Behrens, T.E., and Wallis, J.D. (2011). Double dissociation of value computations in orbitofrontal and anterior cingulate neurons. Nature Neuroscience 14, 1581–1589.

Kennerley, S.W., Dahmubed, A.F., Lara, A.H., and Wallis, J.D. (2009). Neurons in the frontal lobe encode the value of multiple decision variables. Journal of Cognitive Neuroscience 21, 1162–1178.

Levy, D.J., and Glimcher, P.W. (2011). Comparing apples and oranges: Using reward-specific and reward-general subjective value representation in the brain. Journal of Neuroscience 31, 14693–14707.

McDannald, M.A., Esber, G.R., Wegener, M.A., Wied, H.M., Tzu-Lan, L., Stalnaker, T.A., Jones, J.L., Trageser, J., and Schoenbaum, G. (2014). Orbitofrontal neurons acquire responses to ‘valueless’ Pavlovian cues during unblocking. eLIFE 3, e02653.

Padoa-Schioppa, C., and Assad, J.A. (2006). Neurons in orbitofrontal cortex encode economic value. Nature 441, 223–226.

Padoa-Schioppa, C., and Conen, K.E. (2017). Orbitofrontal cortex: a neural circuit for economic decisions. Neuron 96, 736–754.

Plassmann, H., O’Doherty, J., and Rangel, A. (2007). Orbitofrontal cortex encodes willingness to pay in everyday economic transactions. Journal of Neuroscience 27, 9984–9988.

Reber, J., Feinstein, J.S., O’Doherty, J.P., Liljeholm, M., Adolphs, R., and Tranel, D. (2017). Selective impairment of goal-directed decision-making following lesions to the human ventromedial prefrontal cortex. Brain 140, 1743–1756.

Rich, E.L., and Wallis, J.D. (2016). Decoding subjective decisions from orbitofrontal cortex. Nature Neuroscience AOP.

Roesch, M.R., and Olson, C.R. (2004). Neuronal activity related to reward value and motivation in primate frontal cortex. Science 304, 307–310.

Roesch, M.R., Taylor, A.R., and Schoenbaum, G. (2006). Encoding of time-discounted rewards in orbitofrontal cortex is independent of value representation. Neuron 51, 509–520.

Rolls, E.T. (1996). The orbitofrontal cortex. Philosophical Transactions of the Royal Society of London B 351, 1433–1443.

Rolls, E.T., Critchley, H.D., Mason, R., and Wakeman, E.A. (1996). Orbitofrontal cortex neurons: role in olfactory and visual association learning. Journal of Neurophysiology 75, 1970–1981.

Rudebeck, P.H., Mitz, A.R., Chacko, R.V., and Murray, E.A. (2013). Effects of amygdala lesions on reward-value coding in orbital and medial prefrontal cortex. Neuron 80, 1519–1531.

Rudebeck, P.H., and Murray, E.A. (2014). The orbitofrontal oracle: cortical mechanisms for the prediction and evaluation of specific behavioral outcomes. Neuron 84, 1143–1156.

Sadacca, B.F., Wied, H.M., Lopatina, N., Saini, G.K., Nemirovsky, D., and Schoenbaum, G. (2018). Orbitofrontal neurons signal sensory associations underlying model-based inference in a sensory preconditioning task. eLIFE 7, e30373.

Schoenbaum, G., Chiba, A.A., and Gallagher, M. (1998). Orbitofrontal cortex and basolateral amygdala encode expected outcomes during learning. Nature Neuroscience 1, 155–159.

Schoenbaum, G., Chiba, A.A., and Gallagher, M. (1999). Neural encoding in orbitofrontal cortex and basolateral amygdala during olfactory discrimination learning. Journal of Neuroscience 19, 1876-1884.

Schoenbaum, G., and Eichenbaum, H. (1995). Information coding in the rodent prefrontal cortex. I. Single-neuron activity in orbitofrontal cortex compared with that in pyriform cortex. Journal of Neurophysiology 74, 733–750.

Schoenbaum, G., Setlow, B., Saddoris, M.P., and Gallagher, M. (2003). Encoding predicted outcome and acquired value in orbitofrontal cortex during cue sampling depends upon input from basolateral amygdala. Neuron 39, 855–867.

Schoenbaum, G., Takahashi, Y.K., Liu, T.L., and McDannald, M. (2011). Does the orbitofrontal cortex signal value? Annals of the New York Academy of Sciences 1239, 87–99.

Schuck, N.W., Cai, M.B., Wilson, R.C., and Niv, Y. (2016). Human orbitofrontal cortex represents a cognitive map of state space. Neuron 91, 1402–1412.

Stokes, M.G., Kusunoki, M., Sigala, N., Nili, H., Gaffan, D., and Duncan, J. (2013). Dynamic coding for cognitive control in prefrontal cortex. Neuron 78, 364–375.

Thorpe, S.J., Rolls, E.T., and Maddison, S. (1983). The orbitofrontal cortex: neuronal activity in the behaving monkey. Experimental Brain Research 49, 93–115.

Tremblay, L., and Schultz, W. (1999). Relative reward preference in primate orbitofrontal cortex. Nature 398, 704–708.

Wilson, R.C., Takahashi, Y.K., Schoenbaum, G., and Niv, Y. (2014). Orbitofrontal cortex as a cognitive map of task space. Neuron 81, 267–279.

Wimmer, G.E., and Shohamy, D. (2012). Preference by association: How memory mechanisms in the hippocampus bias decisions. Science 338, 270–273.

Yang, T., and Murray, E.A. (2018). Orthogonality of representations of reward magnitude, certainty, and volatility in the macaque orbitofrontal cortex. BioRxIv 365080.

Zhou, J., Jia, C., Feng, Q., Bao, J., and Luo, M. (2015). Prospective coding of dorsal raphe reward signals by the orbitofrontal cortex. Journal of Neuroscience 35, 2717–2730.

