## Supplemental figures for "Rat orbitofrontal ensemble activity contains a multiplexed but value-invariant representation of task structure in an odor sequence task"

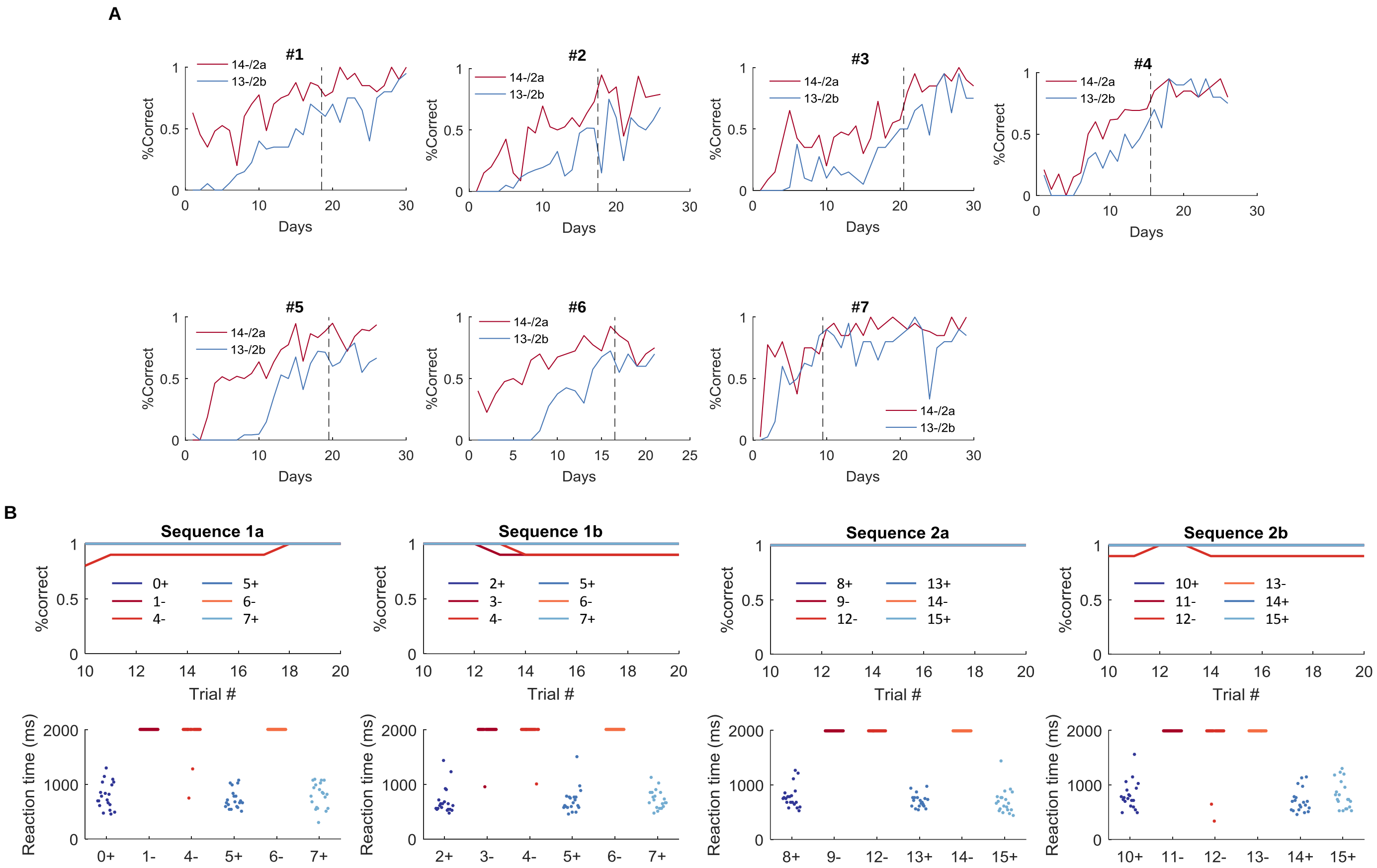


**Figure S1. Behavioral performance during training and recording.** (A) Learning curves on odors 13- and 14- during behavioral training. 7 Rats were trained on S2a and S2b before the full sequence (S1a, S1b, S2a, and S2b). Rats showed gradual increase in behavioral performance (%Correct) on odor 13- in S2b (13-/2b4-) and odor 14- in S2a (14-/2a5-). Plots show learning curves on these two trial types before and after the introduction of the full sequence (The transition is indicated by the vertical dotted lines). Note that rats learned faster on odor 14-/2a5- than odor 13-/2b4-, possibly because rats used reward outcome information on the last trial (13+/2a4+ vs. 13-/2b4-) to facilitate their performance on odor 14-/2a5-, while for performance on 13-/2b4-, rats could only rely on odor information that was two trials back (odor 9-/2a2- vs. odor 11-/2b2-). (B) Behavioral performance on a single session during recording. The performance was assessed by both %Correct and reaction time for reach trial type during the single-unit recording. %Correct is calculated within a ten-trial sliding window. The time window for responding is 2 seconds. Reaction time measures time from leaving the odor port to entry into the water well and is set to be 2 seconds if animals do not respond within the time window.

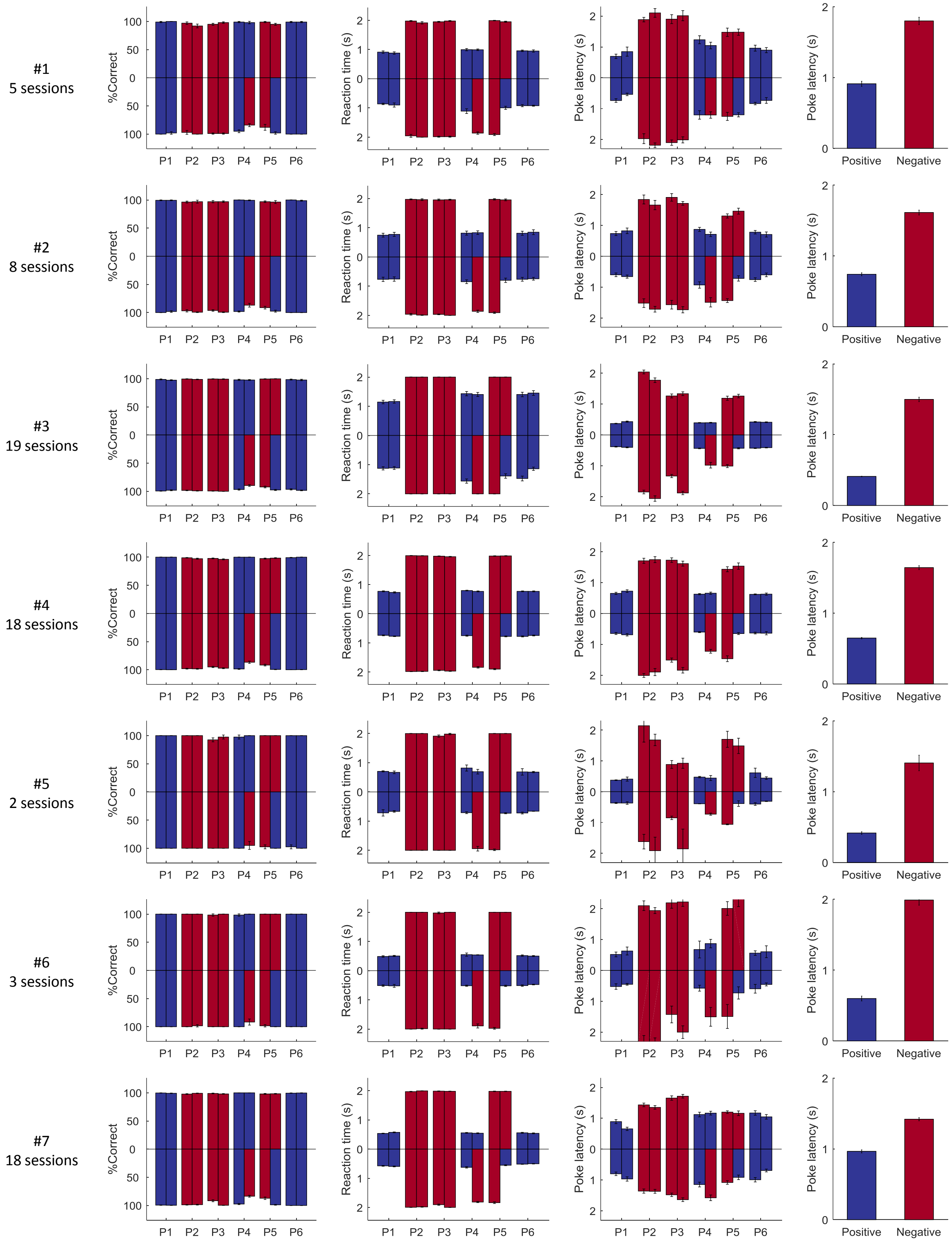


**Figure S2. Behavioral performance of individual rats during recording.**

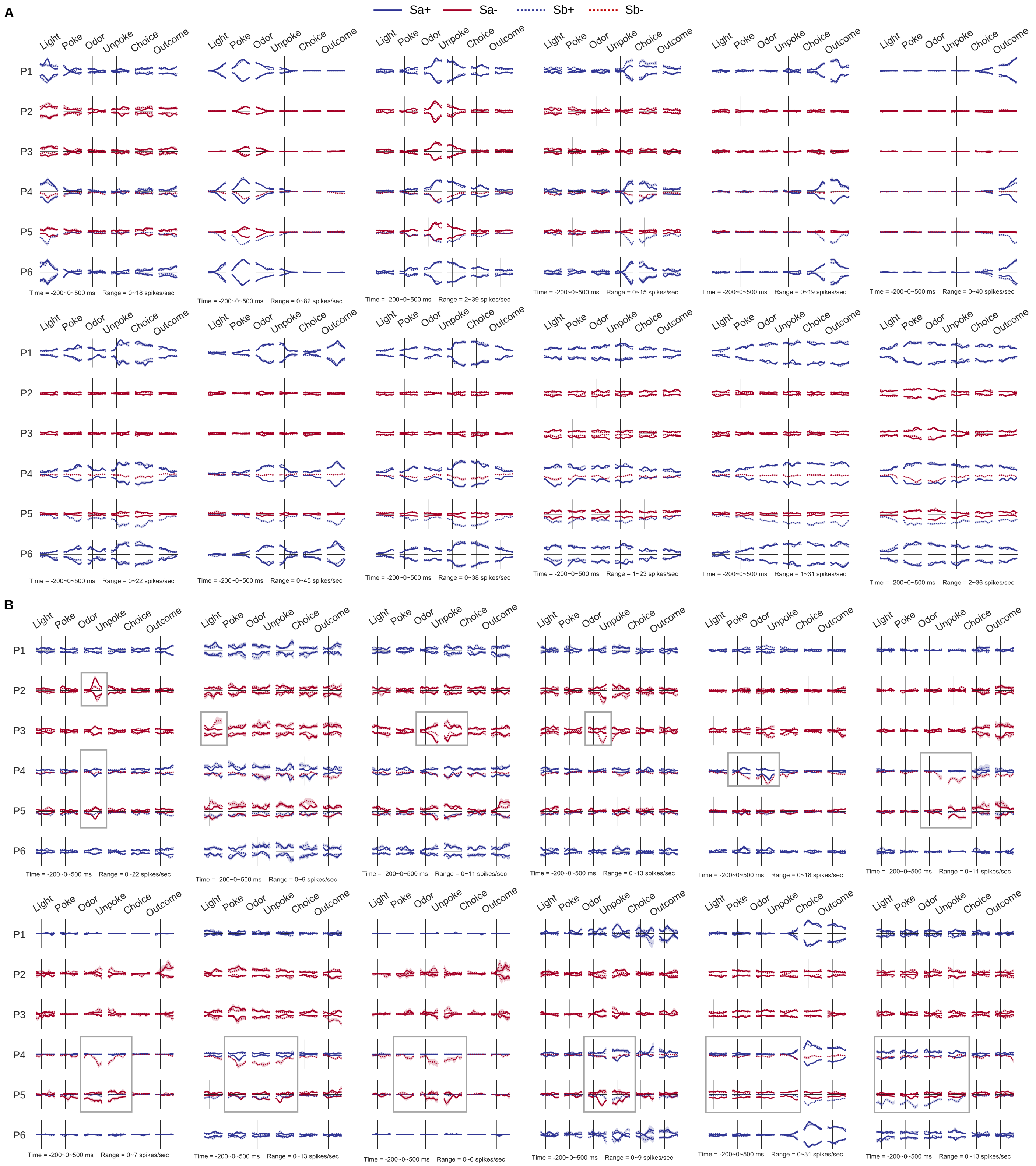


**Figure S3. Activities of example single units.** (A) 12 example single units recorded in the OFC show value-modulated activities. Firing rates for S1 are plotted upwards, while firing rates for S2 are plotted downwards. Note that both of them are non-negative numbers. Dark blue indicates positive trial types, and dark red indicates negative trial types. Solid lines represent Sa, and dashed lines represent Sb. The time and firing rate ranges for each neuron are at the bottom of the plot. The first 6 neurons are transiently activated at only one task event, while the last 6 neurons show more persistent activity throughout several task events. (B) Plots in this figure use the same format as those in **A**. The grey squares highlight the positions where the firing activity cannot be properly explained by value.

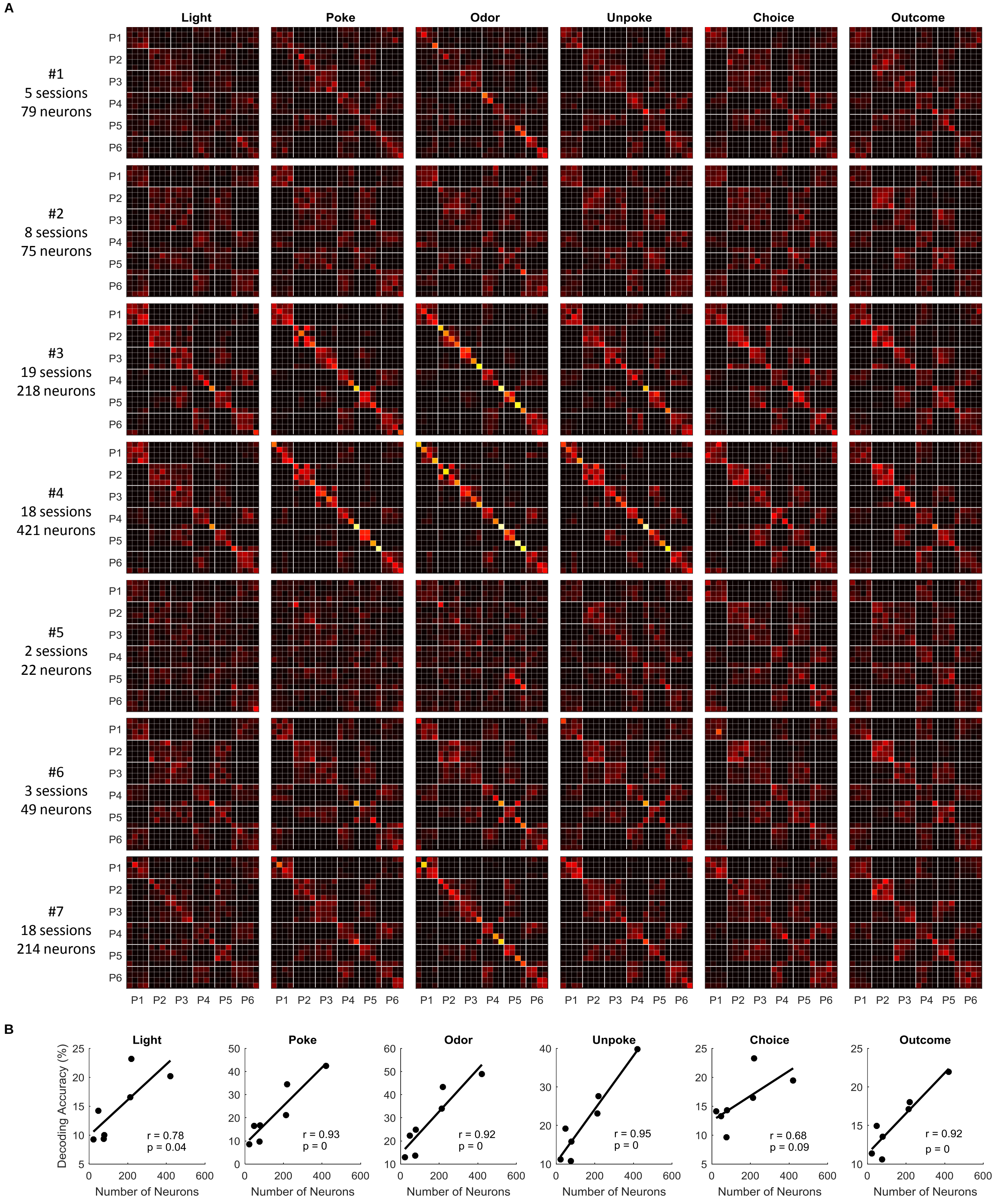


**Figure S4. Decoding of 24 trial types for each rats.** (A) Confusion matrices for each rat (n = 7 rats). (B) Decoding accuracy increases with the number of neurons used in the classifier. Each dot represents the mean accuracy for the decoding of 24 different trial types of each rat. Linear regression analyses show that the mean decoding accuracy and the number of neurons used in the classifier are highly correlated at multiple task events.

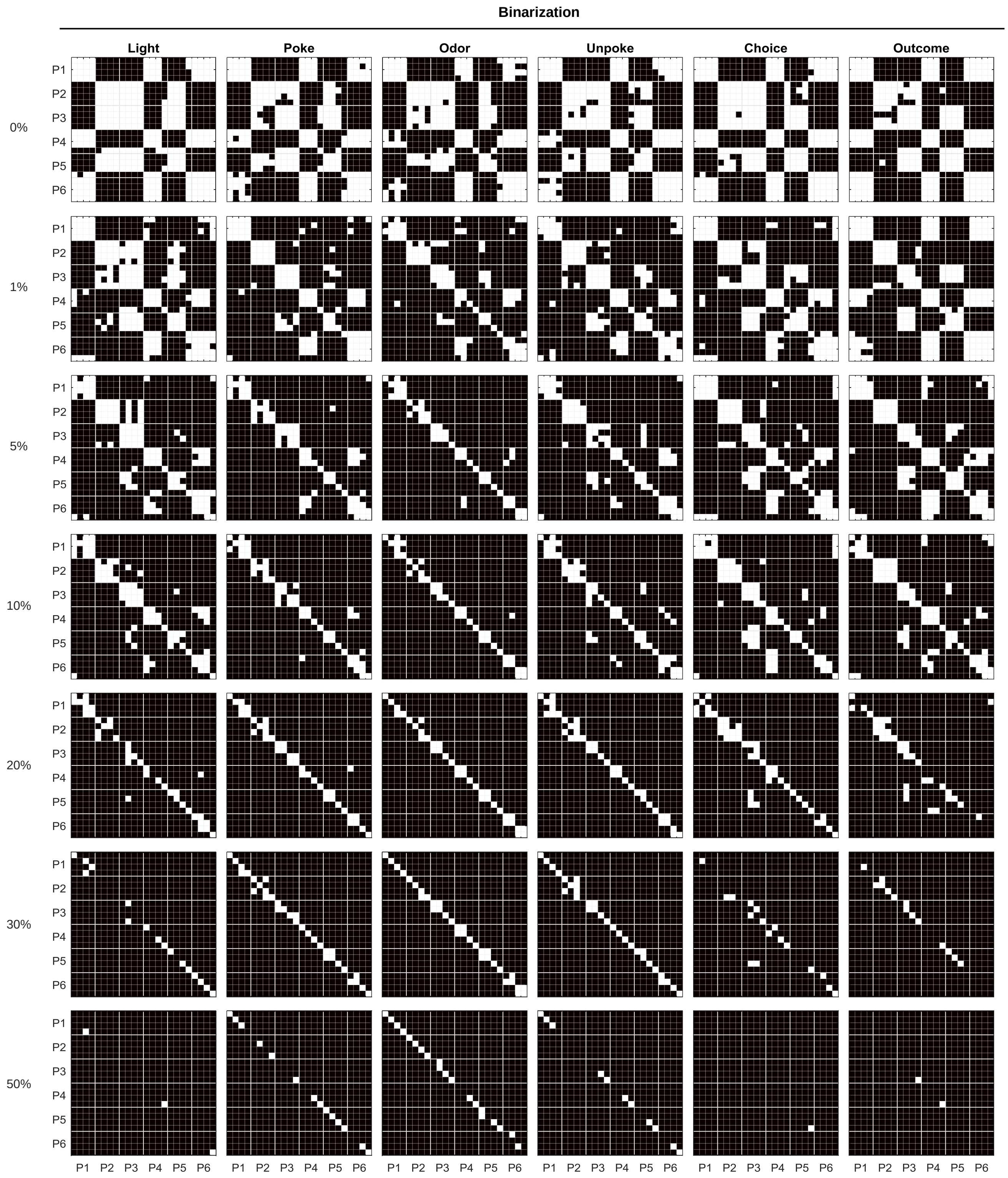


**Figure S5. Binarization of confusion matrices by different thresholds.**
